# Chronic BCR signaling generates and maintains age-associated B cells from anergic B cells

**DOI:** 10.1101/2023.07.26.550463

**Authors:** Keisuke Imabayashi, Yutaro Yada, Miho Ushijima, Motoki Yoshimura, Takeshi Iwasaki, Koichi Akashi, Hiroaki Niiro, Yoshihiro Baba

**Affiliations:** Division of Immunology and Genome Biology, Medical Institute of Bioregulation, Kyushu University, Fukuoka, Japan; Department of Medicine and Biosystemic Science, Kyushu University Graduate School of Medical Sciences, Fukuoka, Japan; Department of Anatomic Pathology, Graduate School of Medical Sciences, Kyushu University, Fukuoka, Japan; Department of Medical Education, Faculty of Medical Sciences, Kyushu University Graduate School of Medical Sciences, Fukuoka, Japan

**Author notes:** Correspondence: Yoshihiro Baba.

## Abstract

Accumulation of age-associated B cells (ABCs) with autoreactive properties contributes to the pathogenesis of autoimmune diseases^1–5^. However, the mechanisms whereby ABCs are generated and maintained are not understood^1, 2, 4^. Here, we show that continuous stimulation of the B-cell receptor (BCR) with self-antigens plays a crucial role in ABC generation from anergic B cells and that this signal is vital for sustaining ABCs during aging and autoimmunity. In ABCs, BCR signaling was constitutively activated and the surface BCR was internalized *in vivo*, as occurs in autoreactive B cells chronically exposed to self-antigens^6^. With aging, ABCs were generated from autoreactive anergic B cells, but not from B cells expressing non-self-reactive BCR. *In vitro* stimulation of anergic B cells with self-antigen, interleukin-21, and Toll-like receptor 7/9 agonists promoted their differentiation to ABCs. Furthermore, the cellular phenotype of ABCs in Bm12-induced lupus mice^7, 8^ resembled that of ABCs in aged mice, showing activation of BCR signaling, expression of activation markers, and BCR internalization. Importantly, Btk was persistently activated in ABCs of aged/autoimmune mice and humans with lupus. Pharmacological Btk inhibition resulted in a marked reduction in the number of ABCs and pathogenicity in lupus mice. Our findings have implications for accumulating ABCs and developing therapies for autoimmune diseases.

## Introduction

Autoreactive B cells play a crucial role in autoimmunity. The body has a system for silencing harmful B cells, called self-tolerance, which can cause peripheral B cells to become anergic^9^. Anergic B cells are induced by chronic exposure of autoreactive B cells to self-antigens via the B-cell receptor (BCR), resulting in a non-responsive state of the BCR and shortened lifespan^6, 10^. Autoreactive B cells accumulate when B cell anergy is disrupted, thereby increasing the risk of developing autoimmune diseases^11–13^. Memory B cells and plasma cells derived from autoreactive B cells are pathogenic in autoimmune diseases^14^; however, a new subset, age-associated B cells (ABCs), has recently been identified as contributing to autoimmunity^1^. ABCs, first reported as a B-cell subset detected in aged mice^2, 3^, accumulate prematurely, independent of aging, in murine lupus models and several human autoimmune diseases, including systemic lupus erythematosus (SLE)^15–19^. However, the mechanisms whereby autoreactive ABCs are generated, maintained, and evade self-tolerance mechanisms are still unknown^1^. Additionally, it remains unclear why ABCs are present in distinct contexts such as aging and autoimmunity^3^. Here, we show that chronic BCR signaling plays a crucial role in the generation of ABCs from anergic B cells through aging, as well as in ABC formation in autoimmunity, ultimately contributing to the progression of autoimmune diseases.

## Results

### BCR signaling is constitutively activated in ABCs

Diverse ABC markers have been used in mice and humans in various contexts including aging, infection, and autoimmunity^4, 20–23^. We sought to clarify ABC markers in aged mice, focusing on their functional aspects. AA4.1^−^CD21^lo^CD23^−^CD11c^+^T-bet^+^ B cells detected in aged mice showed high expression levels of TLR7, TLR9, CD80, and CD86, which correspond to high reactivity to TLR7/9 ligands and strong antigen-presenting capacity to T cells (Extended data Fig. 1a-d). Thus, in this study, we defined the B-cell subset AA4.1^−^CD21^lo^CD23^−^CD11c^+^T-bet^+^ as ABCs. Because ABCs, which are potentially autoreactive B cells, and share antigen specificity with anergic B cells chronically stimulated by self-antigens *in vivo*^6^, we hypothesized that ABCs show chronic self-antigen recognition by the BCR. When we assessed the BCR response on ABCs obtained from aged mice compared with follicular (FO) B cells after anti-IgM stimulation, ABCs showed a less Ca^2+^ increase (Fig. 1a). Notably, ABCs had higher cytosolic Ca^2+^ levels than FO B cells in an unstimulated steady state. These characteristics were similar to those of typical anergic B cells^24^.

**Fig. 1.**
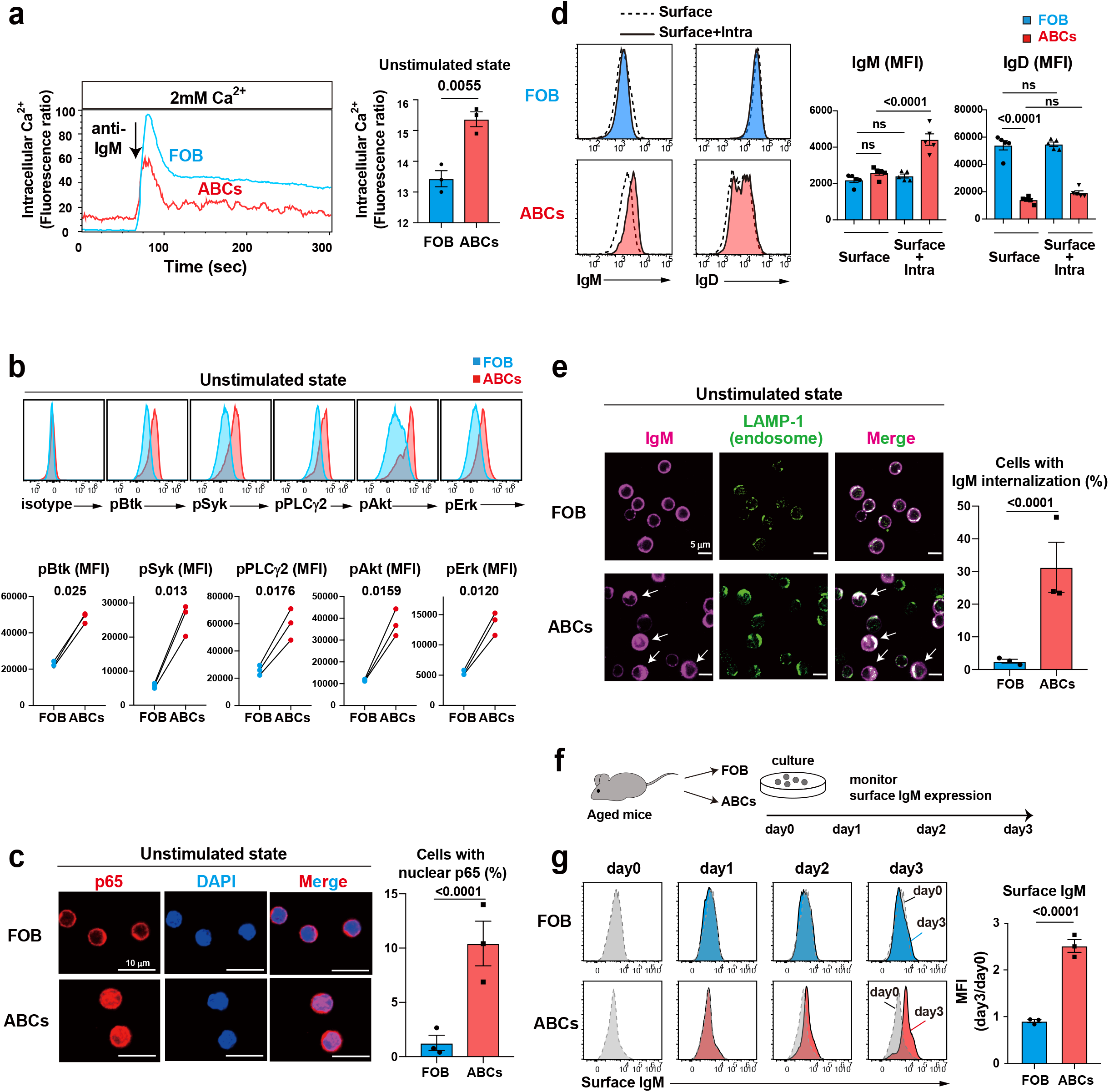
Constitutively active BCR signaling in ABCs via chronic exposure to endogenous antigens. **a**, Ca^2+^-mobilization profiles in splenic FO B cells (AA4.1^−^CD19^+^B220^hi^CD23^+^) and ABCs (AA4.1^−^CD19^+^B220^hi^CD21^lo^CD23^−^CD11c^+^) from aged mice (n=3) (left), and intracellular Ca^2+^ in the unstimulated state (right). **b**, Representative flow cytometry staining (upper) and mean fluorescence intensity (MFI) (lower) of isotype control, pBtk, pSyk, pPLCγ2, pAkt, and pErk of splenic FO B cells and ABCs from aged mice (n=3). **c,** Representative confocal images (left) and the percentage (right) of p65 nuclear translocation in the unstimulated state of FO B cells and ABCs from pooled spleens of aged mice (n=4). Scales bar, 10 μm. **d,** Representative flow cytometry staining (left) and MFI (right) of IgM and IgD gated on splenic FO B cells and ABCs from aged mice (n=3) by surface staining only or surface plus intracellular staining. **e**, Representative confocal images (left) and the percentage (right) of IgM internalization of FO B cells and ABCs from pooled spleens of aged mice (n=4). White arrows indicate internalization of IgM in ABCs. Scales bar, 5 μm. **f**, Schematic of protocol. FO B cells and ABCs from pooled spleens of aged mice (n=4) were left ex-vivo, and surface IgM expression was monitored every 24 h. **g**, Changes of surface IgM levels at day 3 relative to day 0 were compared between FO B cells and ABCs in (g). Data are mean ± s.e.m. *P* values are from two-tailed unpaired **(a,c,e,g)** or paired **(c)** Student’s *t*-test, and two-way ANOVA followed by Tukey’s multiple-comparisons test in **(d)**: ns, not significant. Data are representative of three independent experiments **(a,b,d)**. In the experiment **(c,e,g)**, each dot shows the results of three independent experiments.

Next, we examined the activation status of various intracellular BCR signaling pathways in ABCs. In the unstimulated state, ABCs showed greater phosphorylation of spleen tyrosine kinase (Syk), Bruton’s tyrosine kinase (Btk), phospholipase Cγ2 (PLCγ2), Akt, and Erk than FO B cells (Fig. 1b). In addition, unstimulated ABCs, but not FO B cells, had a robust nuclear translocation of the NF-κB subunit p65 (Fig. 1c). This translocation is not present in anergic B cells^9, 24, 25^. These results show that ABCs have a unique ability to activate the BCR signaling pathway constitutively.

### BCR is internalized in ABCs through continuous antigen recognition

A key question is whether the constitutive activation of BCR signaling in ABCs is due to continuous antigen recognition *in vivo*. When autoreactive B cells are exposed to self-antigens, they can become anergic and internalize their BCR. When crossed MD4 transgenic mice that express BCR with high affinity for hen egg lysozyme (HEL) with ML5 mice expressing soluble HEL in which all B cells were self-specific to the neoself antigen HEL (MD4×ML5), B cells that survived to maturity become anergic^6^. The cell-surface IgM density of these anergic B cells was reduced owing to IgM internalization^25^ (Extended data Fig. 2a). Although we observed that ABCs and FO B cells have comparable levels of surface IgM expression, ABCs exhibited intracellular internalization of IgM (Fig. 1d, Extended data Fig. 2a). Increased surface IgD is a hallmark of anergic B cells^26, 27^; however, ABCs express less surface IgD than FO B cells (Fig. 1d, Extended data Fig. 2a). Confocal microscopy analysis revealed that in FO B cells, internalized IgM was detectable only when the BCR was stimulated (Extended data Fig. 2b), whereas IgM was internalized in LAMP-1-positive endosomes of ABCs in the steady state (Fig. 1e).

**Fig. 2.**
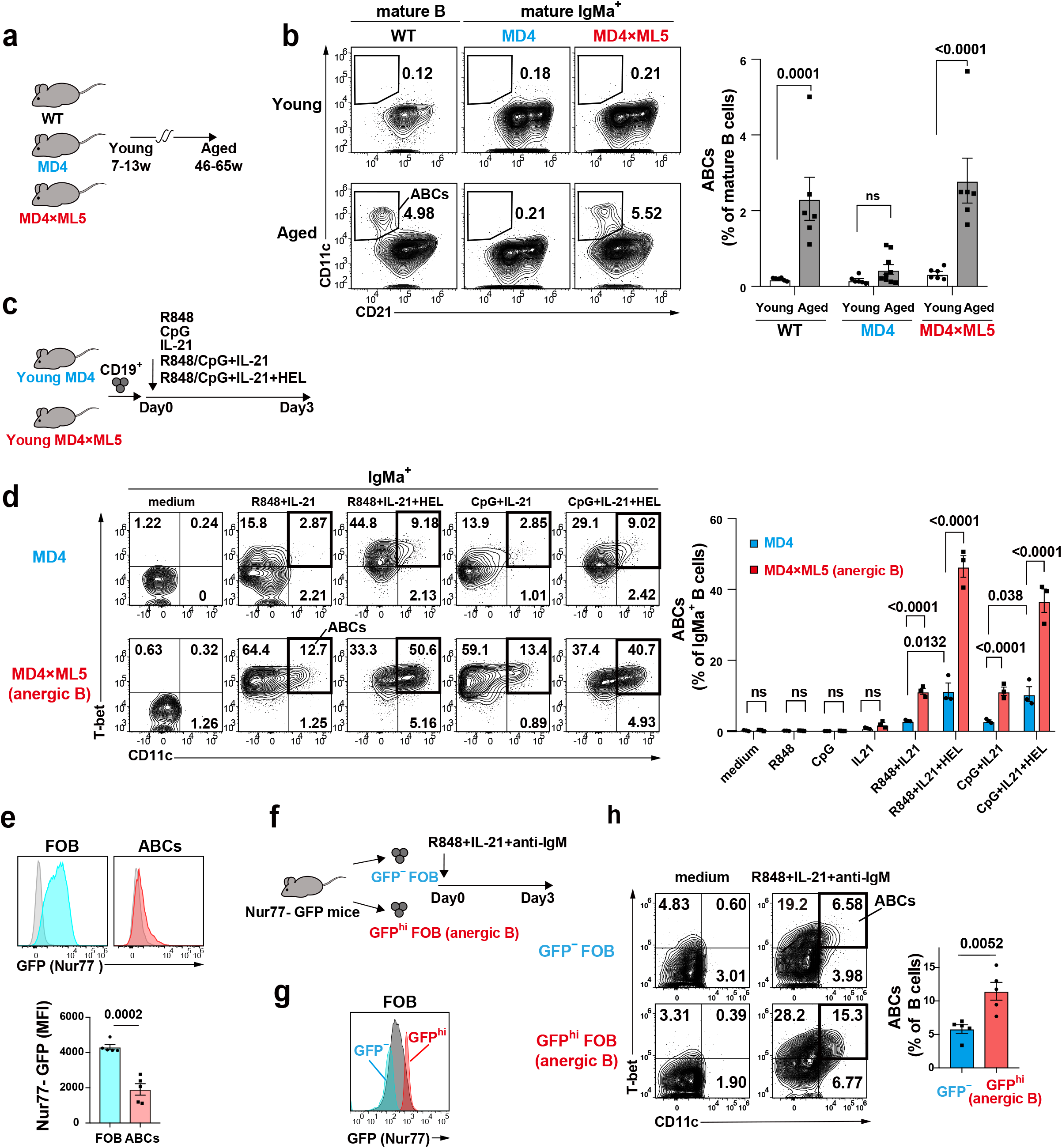
Anergic B cells give rise to ABCs. **a**, Schematic of protocol. Young (7 to 13 weeks) WT mice (n=6), MD4 mice (n=6), MD4×ML5 mice (n=6), and aged (46 to 65 weeks) WT mice (n=6), MD4 mice (n=9), MD4×ML5 mice (n=6) were utilized for analysis. **b**, Representative flow cytometry staining (left) and the percentage (right) of splenic ABCs (CD21^lo^CD11c^+^) among mature B cells (WT) or mature IgMa^+^ B cells (MD4 mice, MD4×ML5 mice) in (a). **c**,. Schematic of protocol. Splenic CD19^+^ B cells were prepared from young MD4 mice (n=3) and MD4×ML5 mice (n=3), and stimulated with cytokines for 3 days. **d**, Representative flow cytometry staining (upper) and the percentage (lower) of CD11c^+^T-bet^+^ ABCs gated on mature IgMa^+^ B cells in (c). **e**, Representative histogram (upper) of Nur77-GFP of splenic FO B cells and ABCs obtained from aged (61-63 weeks) Nur77-GFP mice (n=5). Gray histogram shows GFP expression of each B cells from WT mice. MFI of GFP expression was compared (right). **f**, Schematic of protocol. Splenic GFP^hi^ and GFP^−^ FO B cells were sorted from Nur77-GFP mice (n=5) and stimulated with IL-21, R848, and anti-IgM for 3 days. **g**, Flow cytometry plots of sorted GFP^hi^ (red), GFP^−^ (blue), and overall FO B cells (gray) in (f). **h**, Representative flow cytometry staining (left) and the percentage (right) of ABCs (CD11c^+^T-bet^+^) with or without stimulation in (f). Data are mean ± s.e.m. *P* values are from two-way ANOVA followed by Tukey’s multiple-comparisons test **(b,d)**, and two-tailed unpaired Student’s *t-*test in **(e,h)**: ns, not significant. Data are the pooled data of three independent experiments **(a,b,e-h)**, or representative of three independent experiments **(c,d)**.

Next, we determined whether the IgM internalization is due to continuous antigen recognition *in vivo*. ABCs were placed *ex vivo* to prevent endogenous antigen binding to the BCR, and their surface IgM expression was monitored every 24 hours (Fig. 1f). The surface IgM levels of ABCs, but not of FO B cells, increased over time (Fig. 1g), suggesting that the internalized IgM was recycled to the surface membrane. Surface IgD levels did not change under the same conditions (Extended data Fig. 2c). These findings suggest that ABCs internalize abundant IgM through continuous antigen recognition *in vivo*.

### Anergic B cells give rise to ABCs

To determine whether continuous antigen recognition and constitutive activation of BCR signaling are required for ABC generation, we used MD4 mice, in which almost all mature B cells express a transgenic HEL-specific BCR (IgMa^+^) that limits their response to antigens other than HEL and its homologs^6^. We found that ABCs were generated in aged wild-type mice but not in MD4 mice (Fig. 2a,b), indicating that a diverse BCR repertoire and the recognition of certain endogenous antigens that emerge with age are vital for the development of ABCs. To further explore the *in vivo* requirement of endogenous self-antigens for ABC formation, we analyzed the MD4×ML5 model, in which IgMa^+^ B cells become anergic on chronic stimulation with the self-antigen^6^. The MD4×ML5 mice showed marked ABC generation with age. These findings reveal that ABCs can originate from anergic B cells and highlight the significance of chronic exposure to self-antigens in generating ABCs through aging.

To gain insight into the mechanism of ABC differentiation from anergic B cells, we directly compared naive B cells (MD4 mice) and anergic B cells (MD4×ML5 mice) (Fig. 2c). Compared with MD4×ML5 anergic B cells, MD4 naive B cells differentiated less readily into ABCs when stimulated with TLR7 or TLR9 agonists and interleukin-21 (IL-21), which was promoted by the addition of HEL. In contrast, stimulation of MD4×ML5 anergic B cells with TLR7/9 agonists plus IL-21 was sufficient to produce ABCs from MD4×ML5 anergic B cells, which was further enhanced by HEL stimulation (Fig. 2d). Notably, TLR or IL-21 stimulation alone cannot induce ABCs from MD4×ML5 anergic B cells (Fig. 2d). These results suggest that anergic B cells that were chronically pre-exposed to self-antigens efficiently differentiated into ABCs on TLR and IL-21 stimulation. To further clarify whether anergic B cells give rise to ABCs in normal BCR repertoires, we used Nur77-GFP mice to detect polyclonal anergic B cells. A previous report has shown that GFP^hi^ FO B cells have the characteristics of anergic B cells^28^. We confirmed that GFP^hi^ FO B cells expressed low levels of surface IgM and did not respond to BCR stimulation (Extend Fig. 3a,b). When analyzing aged Nur77-GFP mice, ABCs had lower levels of Nur77 expression (Fig. 2e), which separated them from anergic B cells. We then sorted GFP^hi^ (anergy) and GFP^−^ (non-anergy) FO B cells and stimulated them with TLR7 ligand, IL-21, and anti-IgM (Fig. 2f, g). GFP^hi^ FO B cells differentiated more efficiently into ABCs than GFP^−^ cells (fig. 2h). These results show that anergic B cells induced by self-antigens, whether specific or polyclonal, can differentiate into ABCs.

**Fig. 3.**
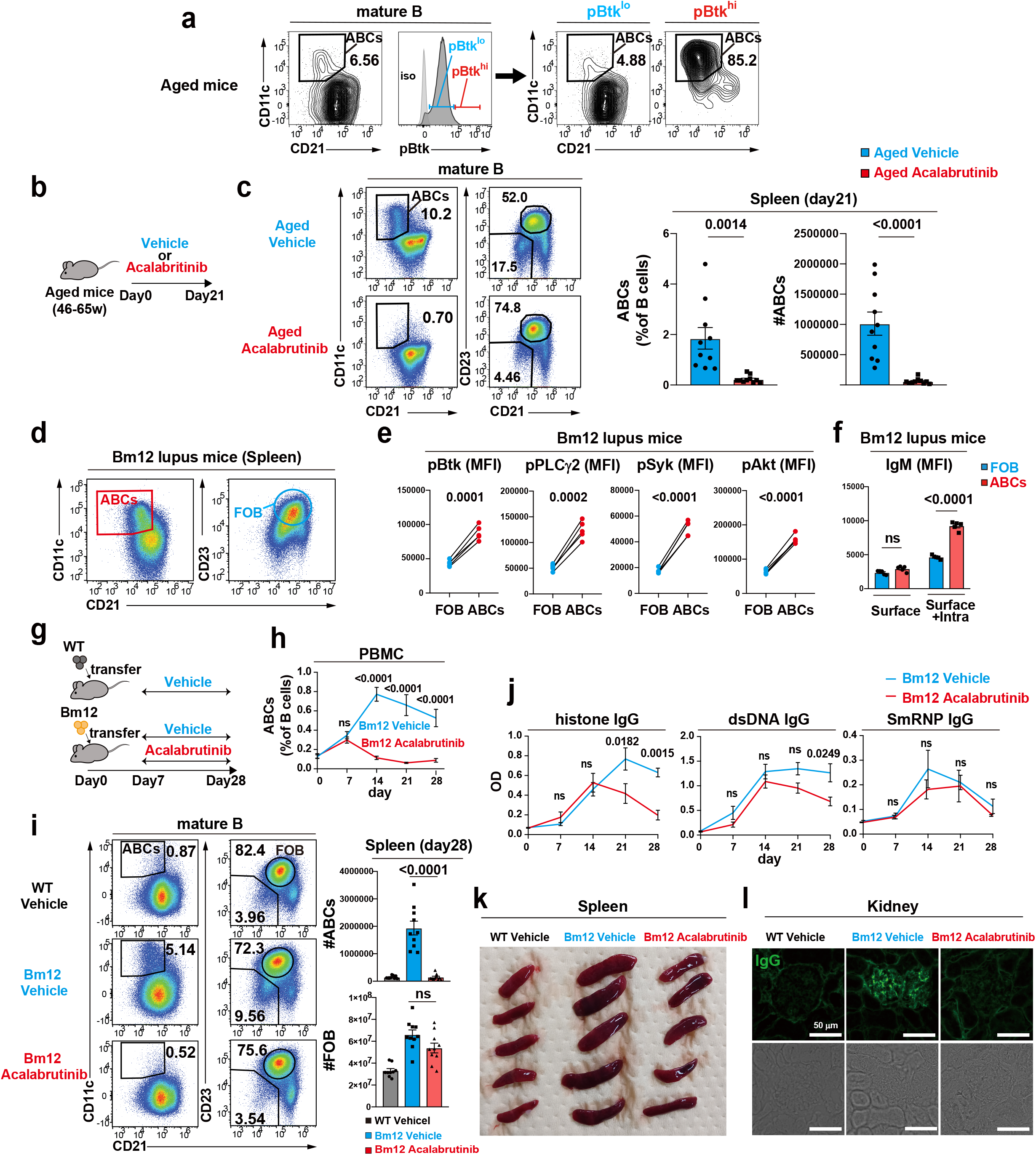
Btk inhibition depletes ABCs and ameliorates autoimmunity in lupus-prone mice. **a**, Flow cytometry plots of splenic ABCs from aged mice (n=3). pBtk^hi^ and pBtk^lo^ mature B cells were gated on CD21 and CD11c. **b**, Aged mice were either given acalabrutinib (n=5, Aged Acalabrutinib) or vehicle (n=5, Aged Vehicle) for 21 days. **c**, Flow cytometry plots (left) and the percentage and absolute number (right) of splenic ABCs in (b). **d**, Representative flow cytometry plots of splenic ABCs and FO B cells obtained from Bm12 lupus mice (n=4). **e**, MFI of pBtk, pSyk, pPLCγ2, pAkt of FO B cells and ABCs from Bm12 lupus mice (n=4). **f**, Calculated MFI of IgM by surface staining only or surface plus intracellular staining of splenic FO B cells or ABCs from Bm12 lupus mice. **g**, Mice started receiving either acalabrutinib (n=5, Bm12 Acalabrutinib) or vehicle (n=5, Bm12 Vehicle) 7 days after Bm12 lymphocytes transfer and were analyzed at day 28. WT mice receiving WT lymphocytes and vehicle (n=5, WT Vehicle) were set as the negative control. **h**, The longitudinal percentage of ABCs among B cells in PBMC in (g). **i**, Flow cytometry plots (left) and absolute number (right) of splenic FO B cells and ABCs in (g). **j**, Autoantibodies against histone, dsDNA, and Sm/RNP in vehicle-or acalabrutinib-treated mice serum in (g). OD, optical density. **k**, Representative spleens from the mice in (g). **l**, Representative renal images of immunoglobulin deposition in kidneys in (g), stained with anti-IgG (upper), and bright field (lower). Scale bars, 50 μm. Data are mean ± s.e.m. *P* values are from two-tailed unpaired **(c)** or paired **(e)** Student’s *t*-test, and one-way **(i)** or two-way **(f,h,j)** ANOVA followed by Tukey’s multiple-comparisons test: ns, not significant. Data are the pool **(c,i)** or representative of two independent experiments **(a-b,d-h,j-l)**.

### Btk is essential for maintenance of ABCs in aged mice

Next, we assessed how B cells persist as an ABC compartment during aging. As Btk, a central component of the BCR downstream pathway, is highly phosphorylated in ABCs (Fig. 3a), we used the Btk inhibitor acalabrutinib to determine whether Btk signaling is required for ABC maintenance in aged mice. Acalabrutinib, a second-generation Btk inhibitor, is highly specific for Btk and has minimal effect on other kinases^29, 30^. Treatment of aged mice with acalabrutinib via drinking water for 3 weeks led to a marked reduction in the number of ABCs (Fig. 3b,c). These data suggest that continuous endogenous antigen recognition via BCR, followed by chronic Btk signaling are critical mechanisms for preserving ABCs.

### ABCs in autoimmune mice exhibit similar characteristics to ABCs in aged mice

To examine whether the features of ABCs in aged mice could also be applied to autoimmune settings, we used a Bm12-induced lupus mouse model^7, 8^. Adoptive transfer of lymphocytes from Bm12 mice to C57BL/6 mice results in the expansion of follicular helper T cells (Tfh), germinal center B cells, plasma cells, and ABCs, leading to autoantibody production and inflammation of organs, including the kidneys, liver, and lungs^8, 18, 31, 32^. We observed that Bm12-induced ABCs exhibited functional markers similar to those in naturally occurring ABCs during aging (Fig. 3d, Extended data Fig. 4a,b). Importantly, Bm12-induced ABCs showed enhanced phosphorylation of Btk, PLCγ2, Syk, and Akt, and BCR internalization in the unstimulated state (Fig. 3e,f). When Bm12 lymphocytes were transferred to Myd88 deficient (Myd88KO) mice, no ABCs were observed (Extended data Fig. 4c). This indicates that, in this model, ABC generation depends on TLR signaling, which is consistent with the increase in ABCs with age^3^. These results suggest that similar factors may operate in aging and autoimmunity to determine the fate of ABC differentiation.

**Fig. 4.**
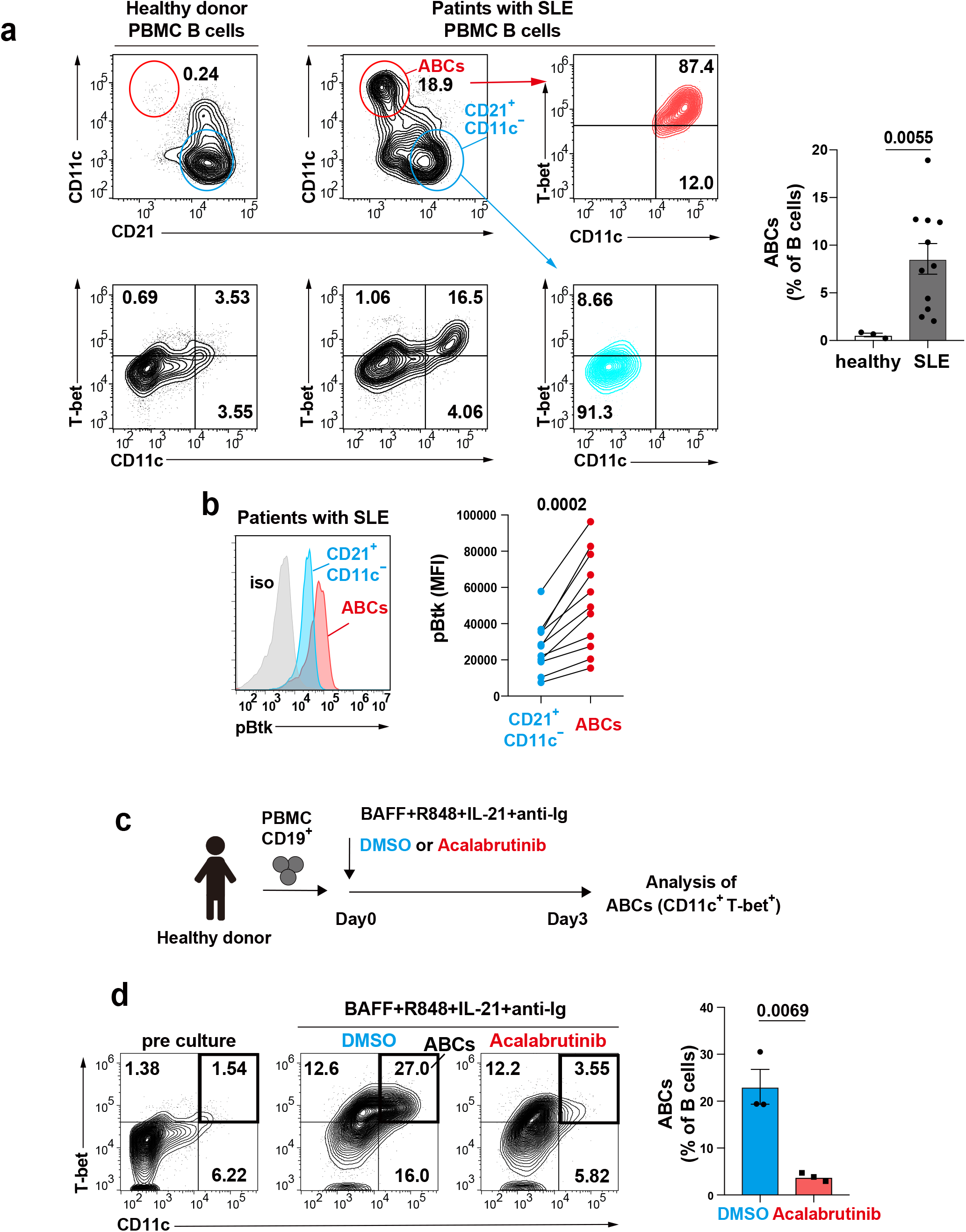
Btk is crucial for the differentiation of human ABCs. **a**, Representative flow cytometry plots of CD21, CD11c, T-bet derived from PBMC (peripheral blood mononuclear cells) of healthy donors (n=3) or SLE (Systematic Lupus Erythematosus) patients (n=11). CD21^lo^CD11c^+^ B cells (ABCs, red) and CD21^+^CD11c^−^ B cells (blue) from PBMC of patients with SLE were gated on the staining of CD11c and T-bet. The percentage of ABCs among B cells was compared. **b**, MFI of pBtk gated on ABCs (red) and CD21^+^CD11c^−^ B cells (blue) derived from PBMC of patients with SLE. **c**, Schematic of protocol. CD19**^+^** B cells collected from PBMC of healthy donors (n=3) were cultured with R848, IL-21, anti-Ig, and BAFF supplemented with Acalabrutinib or DMSO for 3 days, and the percentage of ABCs (CD11c^+^T-bet^+^) was assessed by flow cytometry. **d**, Representative flow cytometry plots and the percentage of ABCs of pre-culture (left), cultured with DMSO (middle, blue) or Acalabrutinib (right, red) in (c). N=3 technical replicates per condition. Data are mean ± s.e.m. *P* values are from two-tailed paired **(b)** or unpaired **(d)** Student’s *t*-test and Mann-Whitney U-test **(a)**. Data are the pooled data of three independent experiments **(a,b)**, and representative of three independent experiments **(c,d)**.

### Btk inhibition depletes ABCs and ameliorates autoimmunity in lupus mice

In autoimmune diseases, ABCs are hypothesized to be pathogenic B cells originating from autoreactive B cells^1, 33^. As constitutive activation of Btk was observed in the ABCs of lupus mice, we investigated whether a Btk inhibitor could remove ABCs and have a therapeutic effect on autoimmune diseases. On day 7 after the transfer of Bm12 lymphocytes, when ABCs and autoantibodies began to be detected (Fig. 3h,j), the mice were given drinking water containing acalabrutinib or vehicle (Fig. 3g). Treatment with acalabrutinib markedly reduced the number of ABCs, but not FO B cells, in the blood and spleen of Bm12-induced lupus mice (Fig. 3h,i). Acalabrutinib also reduced Tfh, Th1, IFNγ^+^CD8^+^T cells, and plasma cells, which can be attributed to the loss of ABCs (Extended data Fig. 5a).

Next, we assessed the effects of acalabrutinib on the autoimmune pathogenesis. Acalabrutinib improves splenomegaly in Bm12-induced mice (Fig. 3k, Extended data Fig. 5b). ABCs detected in the kidneys and liver, the target organs of SLE, decreased after acalabrutinib treatment (Extended data Fig. 5c). Autoantibodies were also reduced after the disappearance of ABCs from peripheral blood (Fig. 3j). Consequently, mice with Bm12-induced lupus treated with acalabrutinib showed a marked decrease in IgG deposits in the kidneys and aggregation of lymphocytes around small vessels in the liver (Fig. 3l, Extended data Fig. 5d). Thus, these results indicate that the formation and/or maintenance of ABCs in autoimmune diseases are dependent on Btk signaling and that its inhibition ameliorates disease progression.

### Btk is crucial for the differentiation of human ABCs

To extend the findings in mouse ABCs to humans, we investigated whether Btk is activated in the ABCs of humans with lupus. As previously described^5^, ABCs with CD21^lo^CD11c^+^T-bet^+^ were expanded in the peripheral blood of patients with SLE (Fig. 4a). ABCs showed higher phosphorylation of Btk than CD21^+^CD11c^-^ B cell populations in the steady state (Fig. 4b). Next, we examined the role of Btk in the generation of human ABCs. When B cells isolated from the peripheral blood of healthy donors were cultured with IL-21, R848 (a TLR7 agonist), anti-Ig, and B cell activating factor belonging to the tumor necrosis factor family (BAFF), they gave rise to ABCs^4, 5, 34^ (Fig. 4c,d). Acalabrutinib treatment was found to inhibit their differentiation (Fig. 4d). Hence, Btk is highly activated under lupus conditions and is required for human ABC differentiation, suggesting that Btk inhibitors could have a therapeutic effect in patients with SLE.

## Discussion

Understanding the process of generation and maintenance of autoreactive ABCs is crucial for elucidating the mechanisms underlying autoimmune diseases, developing diagnostic tools, and identifying therapeutic targets. Here, we show that chronic exposure to endogenous antigens arising from aging and autoimmunity, followed by constitutive activation of BCR signaling, shapes the formation and maintenance of ABCs. We further examined the ontogeny of ABCs and demonstrated that ABCs can originate from anergic B cells.

These findings show that in aged and autoimmune mice, the BCR on ABCs chronically recognizes and incorporates antigens into the cell, which can result in continuous activation of BCR signaling. These results support the previous findings that ABCs have mutated BCR^35^ or characteristics similar to those of memory B cells^20, 23, 36^. We also demonstrated that the development of ABCs depends on self-antigen-mediated B-cell stimulation, as it was abrogated by replacing the polyclonal BCR repertoire with a monoclonal, non-self-reactive BCR in aged mice. Although the expression of TLR7/9, CD40, and major histocompatibility complex (MHC) class II in B cells has been reported to play a role in ABC development^1, 3, 35^, these data reveal that chronic self-antigen-driven BCR signaling is essential for ABC differentiation *in vivo.* Given that chronic exposure to self-antigens causes autoreactive B cells to become anergic, the question arises as to whether ABCs are anergic B cells. ABCs resemble anergic B cells in that they display continuously high levels of cytosolic Ca^2+^ and BCR internalization in the steady state and have a poor BCR response. However, unlike anergic B cells, ABCs retain their BCR reactivity, albeit weakly. A possible explanation is that anergic B cells downregulate surface IgM, whereas ABCs internalize surface IgM, but retain it to the same extent as FO B cells. Decreased surface IgD expression on ABCs may also be involved in retaining BCR reactivity because anergic B cells enable tolerance through increased surface expression of IgD^26, 27, 37^. Moreover, extensively enhanced BCR signaling in ABCs including Syk, Btk, and Akt, and NF-κB is also distinct from that in anergic B cells^38^. Additionally, ABCs are strongly induced to differentiate into plasma cells by TLR7 or TLR9 agonist stimulation, and have a high antigen-presenting capacity to T cells which is not seen in anergic B cells^39^. Given our findings that ABCs can originate from anergic B cells *in vivo* and *in vitro*, it seems likely that ABCs are formed by bypassing B-cell anergy but have inherited some of the characteristics of anergic B cells, although a detailed molecular interplay remains unclear.

Further investigation is required to determine whether ABCs also originate from anergic B cells in autoimmune diseases. Developing therapies that selectively target and suppress the activity of pathogenic autoreactive B cells could minimize immune-related damage while preserving overall immune function. Our data show that targeting BCR signaling via Btk inhibitors reduced ABCs in aged and autoimmune settings, suppressing the pathogenesis of disease. Btk was also highly activated in ABCs of patients with lupus. Therefore, blocking chronic BCR signaling through Btk inhibition in pathogenic ABCs is a potential treatment option for autoimmune diseases.

### Methods Mice

C57BL/6 mice were purchased from CLEA Japan. B6(C)-*H2*-*Ab1^bm^*^12^/KhEgJ (Bm12) and Nur77-GFP transgenic mice were purchased from the Jackson Laboratory. MyD88KO mice were kindly provided by Dr. Shizuo Akira^40^. MD4×ML5 double transgenic were generated by crossing HEL-specific MD4 transgenic mice^6^ to ML5 sHEL transgenic mice^41^. Young mice refer to 7 to 13 weeks unless otherwise noted. Aged mice were bred until the age of 46 to 65 weeks. Mice were bred and maintained under specific pathogen-free conditions. All studies and procedures were approved by the Animal Experiment Committee of Kyushu University. All animal experiments were conducted in accordance with the ARRIVE guidelines and the ethical guidelines of Kyushu University.

### Flow cytometry and antibodies

Single-cell suspensions were stained with the following fluorochrome-conjugated antibodies: For flow cytometry of mouse samples, CD93-BV421 (AA4.1), CD19-BV605 (1D3), CD19-BV650 (1D3), TLR9-PE (J15A7), Fas-BV421 (Jo2), CXCR5-biotin (2G8), CD80-BV650 (16-10A1), Blimp1-PE (6D3), IgMa-biotin (DS-1) phospho-Btk/Itk (pY223/pY180)-PE (N35-86), and streptavidin-BV650 were purchased from BD Biosciences. CXCR4-biotin (2B11), CD43-biotin (eBioR2/60), T-bet-PE (4B10), phospho-PLCγ2-PE (Tyr759) (4NPRN4) were from eBiosciences/Invitrogen. Phospho-Syk (Tyr525/526) (C87C1), phospho-Akt (Ser473) (D9E), phospho-Erk (Thr202/Tyr204) (D13.14.4E), anti-rabbit IgG (H+L)-PE, and isotype control (Rabbit (DA1E) mAb) were from Cell Signaling. B220-APC-Cy7 (RA3-6B2), B220-PE-Cy7 (RA3-6B2), CD3e-FITC (145-2C11), CD4-FITC (GK1.5), CD4-PE-Cy7 (GK1.5), CD5-FITC (53-7.3), CD5-APC (53-7.3), CD8a-PerCP-Cy5.5 (53-6.7), CD11b-PerCP-Cy5.5 (M1/70), CD11c-Alexa647 (N418), CD11c-PE (N418), CD11c-biotin (N418), CD19-FITC (1D3), CD19-BV421 (1D3), CD21/35-PE-Cy7 (7E9), CD21/35-FITC (7E9), CD23-PE (B3B4), CD23-APC (B3B4), CD23-FITC (B3B4), CD40-FITC (HM40-3), CD44-FITC (IM7), CD44-PE-Cy7 (IM7), CD62L-APC (MEL-14), CD69-PE (H1.2F3), CD73-PE (TY/11.8), CD80-PE (16-10A1), CD86-APC (GL-1), CD138-BV421 (281-2), CXCR3-BV650 (CXCR3-173), I-A/I-E-APC (M5/114.15.2), ICOSL-biotin (HK5.3), IFN-γ-FITC (XMG1.2), IgM-FITC (RMM-1), IgMa-biotin (MA-69), IL-17A-APC (TC11-18H10.1), PDL2-biotin (TY25), PD1-APC-Cy7 (29F.1A12), T-bet-Alexa647 (4B10), TCRβ-BV421 (H57-597), TCRβ-APC-Cy7 (H57-597), TLR7-PE (A94B10), streptavidin-APC, streptavidin-PerCP-Cy5.5, and streptavidin-PE were from BioLegend. Zombie aqua or zombie NIR dye (BioLegend) was used for detecting dead cells. For flow cytometry of human samples, CD21-FITC (Bu32), CD27-APC (M-T271), CD19-APC-Cy7 (HIB19), CD11c-BV421 (S-HCL-3), and IgD-PE-Cy7 (IA6-2) were from BioLegend. All antibodies were titrated prior to use to determine optimal working dilutions.

For flow cytometry, mouse splenocytes were red blood cell lysis with ammonium chloride/potassium chloride (ACK) solution. Cells were washed and suspended with FACS staining buffer (PBS (Nacalai Tesque) containing 2% (vol/vol) FCS and 1 mM EDTA) and blocked with unconjugated anti-CD16/32 (2.4G2, Tombo) in the staining buffer for 15 min, followed by staining with surface antibody cocktails at 4℃ for 20 min.

For the detection of intracellular cytokine, cells were first stimulated with 500 ng/ml ionomycin and 50 ng/ml PMA in the presence of monensin (Invitrogen) for 4 h at 37℃ with 5% CO_2_ in RPMI complete medium (RPMI containing 10% FCS, HEPES, L-Glutamine, penicillin/streptomycin, sodium pyruvate (Nacalai Tesque)).

For intracellular staining, cells were fixed and permeabilized with IC fixation buffer (eBioscience) or Foxp3 Staining Buffer Set (eBioscience) for 20 min and washed with permeabilization buffer (eBioscience) before incubation with anti-TLR7, anti-TLR9, anti-IFN-γ, anti-IL-17A, anti-T-bet, or anti-Blimp1 antibody.

For the detection of phospho-Btk (pBtk), phospho-Syk (pSyk), phospho-PLCγ2 (pPLCγ2), phospho-Akt (pAkt), and phospho-Erk (pErk), cells were first stained with surface antibody cocktails and then incubated at 37 °C for 10 min in a volume of 500 μL RPMI complete medium, and immediately fixed with the same volume of BD Phosflow^TM^ Fix Buffer I (BD Biosciences) at 37 °C for 10 min. Cells were then permeabilized with ice-cold BD Phosflow^TM^ Perm Buffer II (BD Biosciences) for 30 min on ice, followed by staining with anti-pBtk, anti-pSyk, anti-pPLCγ2, anti-pAkt, and anti-pErk following the manufactures’ instructions. Data were collected on Cytoflex (Beckman Coulter) and analyzed in FlowJo (TreeStar).

For B cell sorting, splenic B cells were enriched by a positive selection of CD19^+^ cells with anti-CD19 magnetic beads (Miltenyi Biotec). Enriched B cells were stained by surface antibody cocktails and further sorted by gating FO B cells (CD19^+^B220^hi^AA4.1^−^CD23^+^), ABCs (CD19^+^B220^hi^AA4.1^−^CD21^lo^CD23^−^CD11c^+^), or CD21^lo^CD23^−^CD11c^−^ B cells (CD19^+^B220^hi^AA4.1^-^CD21^lo^CD23^−^CD11c^−^) using FACSMelody (BD Biosciences).

### *In vitro* culture assay

For *in vitro* culture of B cells from aged mice, sorted B cells were washed and resuspended in RPMI complete medium (RPMI containing 10% FCS, HEPES, L-Glutamine, penicillin/streptomycin, sodium pyruvate). 5×10^4^ B cells were cultured per well in 96 well round plates containing media alone or media containing 0.5 μg/ml R848 (InvivoGen) or 5.0 μg/ml CpG (InvivoGen) at 37℃ with 5% CO_2_ for 3 days.

For co-culture of B cells and T cells, splenic OT-II transgenic naïve CD4^+^ T cells (TCRβ^+^CD4^+^CD44^lo^CD62L^hi^CD25^−^) were sorted using FACSMelody. B cells were co-cultured with 5×10^4^ CTV (Cell Trace Violet)-labeled CD4^+^ T cells at a 1:1 ratio in the presence of 0.4 μg/ml OVA_323-339_ peptide (MBL) for 3 days.

For *in vitro* induction of ABCs from MD4 mice and MD4×ML5 mice, splenic B cells isolated by positive selection with anti-CD19 magnetic beads (>95% positive for B220 staining) were cultured in the presence of the appropriate combination of the following; 0.5 μg/ml R848, 5.0 μg/ml CpG, 30 ng/ml mIL-21(R&D), and 5.0 μg/ml HEL protein for 3 days.

For *in vitro* induction of ABCs from Nur77-GFP mice, FACS-sorted GFP^hi^ and GFP−FO B cells were cultured in the presence of 0.5 μg/ml R848, 30 ng/ml mIL-21, 1.0 μg/ml anti-IgM F(ab’)_2_ (Jackson ImmunoResearch Laboratories) for 3 days.

For *in vitro* induction of ABCs from human PBMCs, human PBMCs were isolated by density gradient centrifugation using Lympholyte (Cedarlane) or Lymphoprep (Axis-Shield). B cells were purified by using positive selection with anti-CD19 magnetic beads (>95% positive for CD20 staining) and were cultured in the presence of 20 ng/ml hBAFF (R&D systems), 1 μg/ml R848 (Invitrogen), 50 ng/ml hIL-21 (BioLegend), and 10 μg/ml goat anti-human IgA+IgG+IgM (H+L) (Jackson ImmunoResearch Laboratories) supplemented with 1 μM acalabrutinib (MedChem Express) or DMSO for 3 days.

### Assessment of IgM/IgD internalization and re-expression by flow cytometry

For the assessment of IgM/IgD internalization by flow cytometry, splenocytes were stained with surface antibody cocktails (anti-CD11c, anti-B220, anti-AA4.1, anti-CD19, anti-CD23, and anti-CD21) and then fixed with IC fixation buffer for 20 min at 4℃. For the detection of surface IgM/IgD, cells were washed with PBS and stained with anti-IgM and anti-IgD in FACS staining buffer at 4℃ for 20 min. For the detection of surface plus intracellular IgM/IgD simultaneously, cells were washed with permeabilization buffer and stained with anti-IgM and anti-IgD diluted in permeabilization buffer at the same concentration of that used in the surface staining at 4℃ for 20 min.

For the assessment of IgM/IgD re-expression, FACS-sorted splenic FO B cells or ABCs from pooled spleens of aged mice were washed and resuspended in RPMI complete medium. 5×10^4^ B cells suspended in 500 μl complete medium were cultured per well in 48 well plates containing 50 ng/ml BAFF (R&D) for survival aid, and their surface IgM/IgD expression was assessed by flow cytometry every 24 h. The mean fluorescence intensity of IgM and IgD was compared between each time point.

### Calcium measurement

For cytosolic calcium measurement, splenocytes were loaded with Cal520 acetoxymethyl ester (Cal520-AM) (AAT Bioqwest), FuraRed-AM (Invitrogen), and Pluronic F127 (Invitrogen) in DMEM containing 2% FCS at 37℃ for 45 min. Cells were washed twice with Ringer solution (155 mM NaCl, 4.5 mM KCl, 1 mM MgCl_2_, 2 mM CaCl_2_, 5 mM Hepes, 5.6 mM D-glucose, 0.025% BSA) and stained with antibodies (anti-B220, anti-CD21, anti-CD23, and anti-CD11c). Steady-state intracellular calcium was measured for 60 sec, and then cells were stimulated with 10 μg/ml anti-IgM F (ab’)_2_ for 5 min. Changes in fluorescence intensity were monitored on FACS Fortessa (Becton Dickinson), and Ca^2+^mobilization was measured by calculating changes in the fluorescence ratio of Cal520 to FuraRed.

### Immunofluorescence analysis of nuclear translocation of p65 and IgM internalization

For the assessment of p65 nuclear translocation, FACS sorted 1×10^5^ FO B cells or ABCs were immediately adhered to cover glasses coated with Cell-Tak (Discovery Labware Inc). Cells were fixed and permeabilized with ice-cold methanol, blocked with Blocking One Hist (Nacalai Tesque) and stained with rabbit anti-p65 (Cell Signaling) at 4℃ overnight. Cells were then stained with Alexa647 conjugated anti-rabbit IgG (H+L) and DAPI (BioLegend) at room temperature for 1 h, protected from light.

For the detection of IgM internalization, cells were immediately adhered to cover glasses or pre-stained with biotin-conjugated anti-IgM F (ab’)_2_ on ice for 30 min and then for 30 min at 37℃ before stacking to cover glasses. Cells were fixed and permeabilized with ice-cold methanol, blocked with Blocking One Hist, and stained with biotin-conjugated goat anti-IgM F (ab’)_2_ (Southern Biotech) and Alexa488 labeled anti-LAMP-1 (Santa Cruz) at 4℃ overnight. Cells were stained with Alexa594 labeled streptavidin and DAPI at room temperature for 1 h. Stained cells were washed in PBS and mounted in ProLong Diamond Antifade (Invitrogen) at room temperature for 24 h. Images were obtained with LSM 700 or LSM 900 confocal microscopy (Zeiss). The number of cells with p65 nuclear translocation and IgM internalization was evaluated by collecting at least 100 cells from randomly taken photographs in each experiment.

### The Bm12-Inducible Model of Systemic Lupus Erythematosus

The Bm12 transfer model, as originally described by Morris et al.^7^ and modified by Klarquist and Janssen^8^ was adopted. CD4^+^ T cells from mice expressing the MHC-II mutant allele I-Abm12 (bm12 mice), which differs from I-Ab by three amino acids, recognize and activate any B cell expressing I-Ab, as evidenced by the production of anti-DNA autoantibodies and lupus phenotypes in host mice adoptively transferred with bm12 lymphocytes^7, 8^. Single-cell suspensions of spleens and lymph nodes (superficial cervical, axillary, mesenteric, and inguinal) obtained from Bm12 donor mice were prepared and resuspended in RPMI-1640 medium containing 10% FCS at room temperature without red blood cell lysis treatment. Lymphocytes were resuspended in PBS, and 3×10^6^ cells were injected intravenously. Donor and recipient mice were age- and sex-matched. For analyzing Myd88 KO mice, 6×10^7^ lymphocytes from Bm12 or WT mice were intravenously transferred to WT or Myd88KO mice and sacrificed for analysis at day 21. For acalabrutinib treatment, 0.15 mg/ml acalabrutinib diluted in sterile water containing 2% hydroxypropyl-β-cyclodextrin (Wako) as a vehicle was prepared. Mice were allowed to either vehicle or acalabrutinib in the drinking water ad libitum 1 week after lymphocyte transfer and exposed to solutions until the end of the experiment for a total of 3 weeks. Serum and PBMC were obtained before and every week after the transfer. Mice were sacrificed for analysis 4 weeks after cell transfer, and spleens, liver, and kidney were harvested and weighed. Single-cell suspension of each organ was prepared for flow cytometry analysis. For preparing a single-cell suspension of kidneys, isolated kidneys were minced finely and treated with 0.5 mg/ml collagenase D (Roche) and 25 unit/ml DNaseⅠ (Worthington) in DMEM containing 10% FCS for 40 min at 37℃, with gentle stirring. Digested kidneys were manually ground using EASY strainer (Greiner Bio-One). Mononuclear cells were isolated using Percoll (Cytiva) density gradient centrifuge at 850 g for 20 min. For preparing single-cell suspension of livers, harvested livers were manually ground using EASY strainer, and mononuclear cells were isolated using Percoll. Single-cell suspensions of both kidneys and livers were obtained after red blood cell lysis using ACK solution and used for flow cytometry analysis.

### Histological analysis

For immunofluorescence studies of mouse kidneys, harvested kidneys were cut along the short axis at the maximum area of the whole kidney, fixed with 4% paraformaldehyde at 4℃ for 2 h, and incubated in 20% sucrose in PBS at 4℃ overnight. OCT-embedded (Sakura) kidneys were cryosectioned into 10 μm sections and mounted on slide glasses. The slides were dried and washed with PBS, followed by blocking with 1% goat serum and 3% BSA in PBS at room temperature for 1 h. The sections were stained with Alexa488 labeled anti-IgG (polyclonal, BioLegend) at 4℃ overnight. For Hematoxylin and Eosin staining (H&E) staining of the liver, mouse livers were prepared, and the right lobe was fixed in 10% formaldehyde, embedded in paraffin, sectioned (4.0 μm thickness), and then stained with H&E. Images of kidneys and livers were obtained with Keyence BZ-X700 microscope (Keyence Corporation).

### ELISA

Plates were coated with 10 μg/ml dsDNA (Invitrogen), 10 μg/ml Histon (Roche), or 0.1 μg/ml Sm/RNP (GenWay) in PBS at 4℃ overnight. The serum was diluted and incubated on coated plates at 4℃ overnight. Plates were then incubated for 3 h with horseradish peroxidase–labeled goat anti-mouse IgG (Southern Biotech). OD_450_ was measured on a microplate reader (iMArkTM Microplate Reader, Bio-Rad).

### Human participants

The study recruited 4 male healthy donors aged 31-36 years with no medication and 11 female patients with SLE attending Kyushu University Hospital aged 22–31 years with mild-to-moderate disease activity. Informed consent was obtained from all patients in accordance with the Declaration of Helsinki. The Institutional Review Board of Kyushu University Hospital approved all research on human subjects (No. 2023-51).

### Statistical analysis

We performed statistical evaluation using Prism software (GraphPad, La Jolla, CA). A two-tailed, unpaired, or paired Student’s *t*-test was applied for the statistical comparison of two groups. In case of unequal variance, the *t*-test with Welch’s correction was used. Comparisons of two nonparametric datasets were made by the Mann-Whitney U-test. Analysis of variance (ANOVA) followed by Tukey’s multiple-comparisons test was applied for statistical comparison between multiple groups. *P* values of less than 0.05 were considered statistically significant.

## Acknowledgments

We thank A. Baba, C. Ono, K. Kawata, and S. Hatano for technical assistance; M. Tanaka and K. Kageyama for technical support in the Medical Institute of Bioregulation, Kyushu University for the animal facility; S. Sawa, Y. Fukui, and T. Kurosaki for important mouse strains; N. Furuno for secretarial help; technical assistance from The Research Support Center, Research Center for Human Disease Modeling, Kyushu University Graduate School of Medical Sciences. We thank editage Group (https://www.editage.com) for editing a draft of this manuscript. This work was partially supported by JSPS KAKENHI (JP18H02626, 19K22537, JP21H02753 to Y.B, and JP21K08460 to H.N.), Agency for Medical Research and Development (AMED) (JP19ek0410044 and JP19gm6110004 to Y.B.), the Medical Research Center Initiative for High Depth Omics and RIIT (to Y.B).

## Author contributions

K.I. performed experiments, analyzed data, and wrote the manuscript. Y.Y. and M.U. provided technical supports of experiments and analyzing data. T.I. provided pathological assessments. M.Y., K.A., and H.N. designed the human study and provided important inputs. Y.B. designed and supervised the study and wrote the manuscript. All authors reviewed and commented on the manuscript.

