## Extended DATA for "Chronic BCR signaling generates and maintains age-associated B cells from anergic B cells"

### Extended data Fig. 1

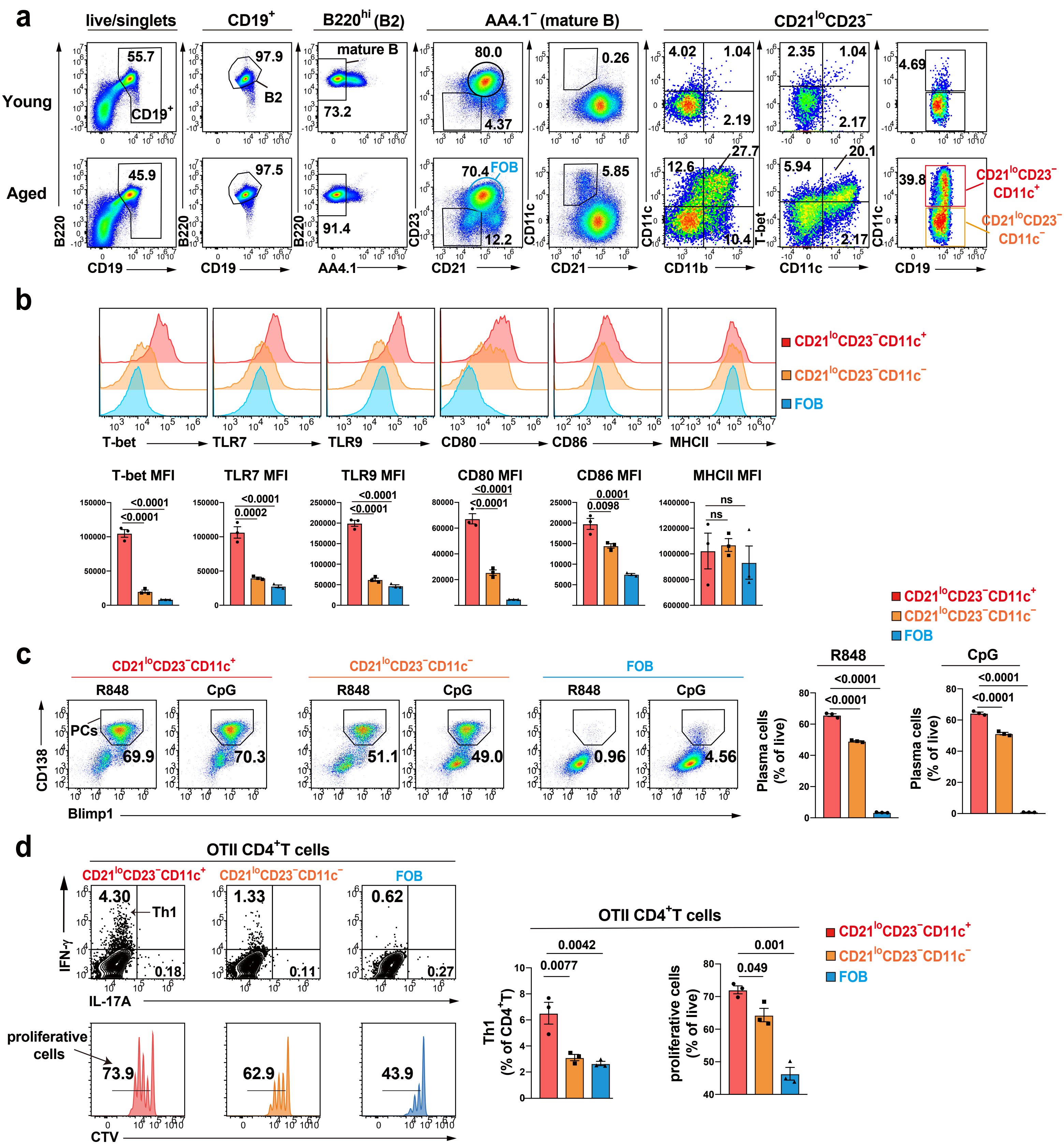


**Extended data Fig. 1 | ABCs are hyper-responsive to TLR7/9 ligand and have high antigen presenting capacity. a,** Representative flow cytometry plots of CD19, B220, AA4.1, CD21, CD23, CD11c, and T-bet of splenocytes from young (n=3) and aged mice (n=3). **b,** Representative histograms and MFI of T-bet, TLR7, TLR9, CD80, CD86, MHCII of splenocytes from aged mice (n=3), indicating CD19$^{+}$B220$^{hi}$AA4.1$^{-}$CD21$^{lo}$CD23$^{-}$CD11c$^{+}$ (CD21$^{lo}$CD23$^{-}$CD11c$^{+}$ B cells, red), CD19$^{+}$B220$^{hi}$AA4.1$^{-}$CD21$^{lo}$CD23$^{-}$CD11c$^{-}$ (CD21$^{lo}$CD23$^{-}$CD11c$^{-}$ B cells, orange), and CD19$^{+}$B220$^{hi}$AA4.1$^{-}$CD23$^{+}$(FO B cells, blue). **c,** FACS sorted CD21$^{lo}$CD23$^{-}$CD11c$^{+}$ B cells, CD21$^{lo}$CD23$^{-}$CD11c$^{-}$ B cells, and FO B cells from pooled splenocytes of aged mice (n=4) were cultured for 3 days with either CpG (TLR9 ligand) or R848 (TLR7 ligand). Blimp1$^{+}$CD138$^{+}$ cells are shown as plasma cells (PCs). N=3 technical replicates per condition. **d,** Flow cytometry staining of CTV-labelled splenic OTII naive CD4$^{+}$ T cells (CD44$^{lo}$CD62L$^{hi}$CD25$^{-}$TCRβ$^{+}$CD4$^{+}$) that were cocultured with splenic CD21$^{lo}$CD23$^{-}$CD11c$^{+}$ B cells, CD21$^{lo}$CD23$^{-}$CD11c$^{-}$ B cells, or FO B cells in the presence of OVA peptide for 3 days, re-stimulated and analyzed for the expression of IFN-γ and IL-17A. IFN-γ$^{+}$ T cells are shown as Th1. Proliferative cells means CTV diluted CD4$^{+}$ T cells. N=3 technical replicates per condition. Data are mean ± s.e.m. *P* values are from one-way ANOVA followed by Tukey's multiple-comparisons test **(b,c,d)**: ns, not significant. Data are representative of two **(c,d)** or three **(a,b)** independent experiments.

### Extended data Fig. 2

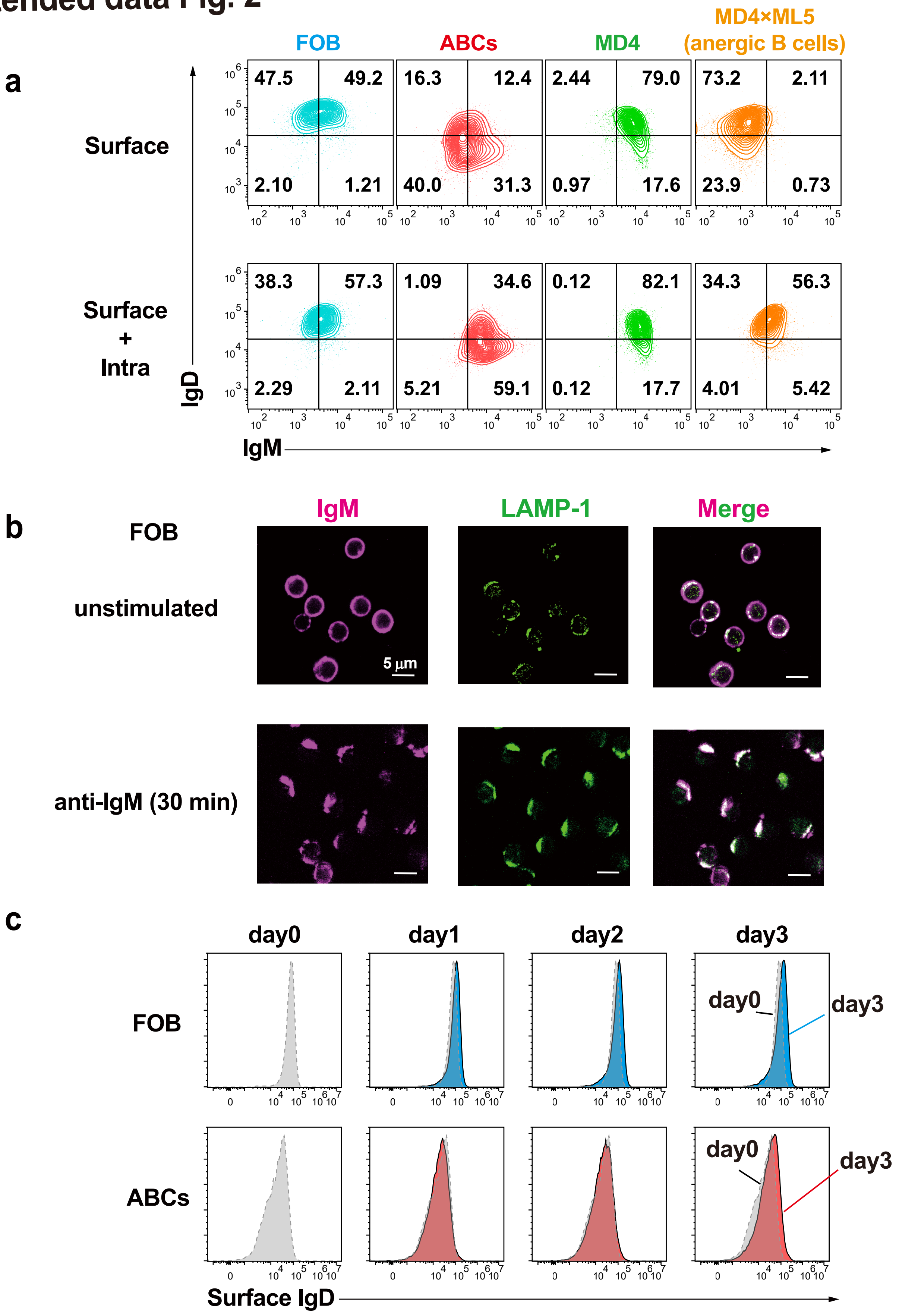


**Extended data Fig. 2 | Surface and intracellular BCR in B cells. a**, Representative surface staining or surface plus intracellular staining of IgM and IgD of splenic FO B cells and ABCs from aged mice (n=3), CD19$^{+}$B cells from young MD4 mice (n=3) and young MD4×ML5 mice (n=3). **b**, Staining of IgM (magenta) and LAMP-1 (green) of sorted FO B cells from pooled splenocytes of aged mice (n=4) with (lower) or without (upper) stimulation with anti-IgM for 30 min assessed by confocal microscopy. Scales bar shows 5 μm. **c**, FO B cells and ABCs from pooled splenocytes of aged mice (n=4) were left ex-vivo, and surface IgD expression was assessed every 24h. Data are representative of two independent experiments **(a-c)**.

### Extended data Fig. 3

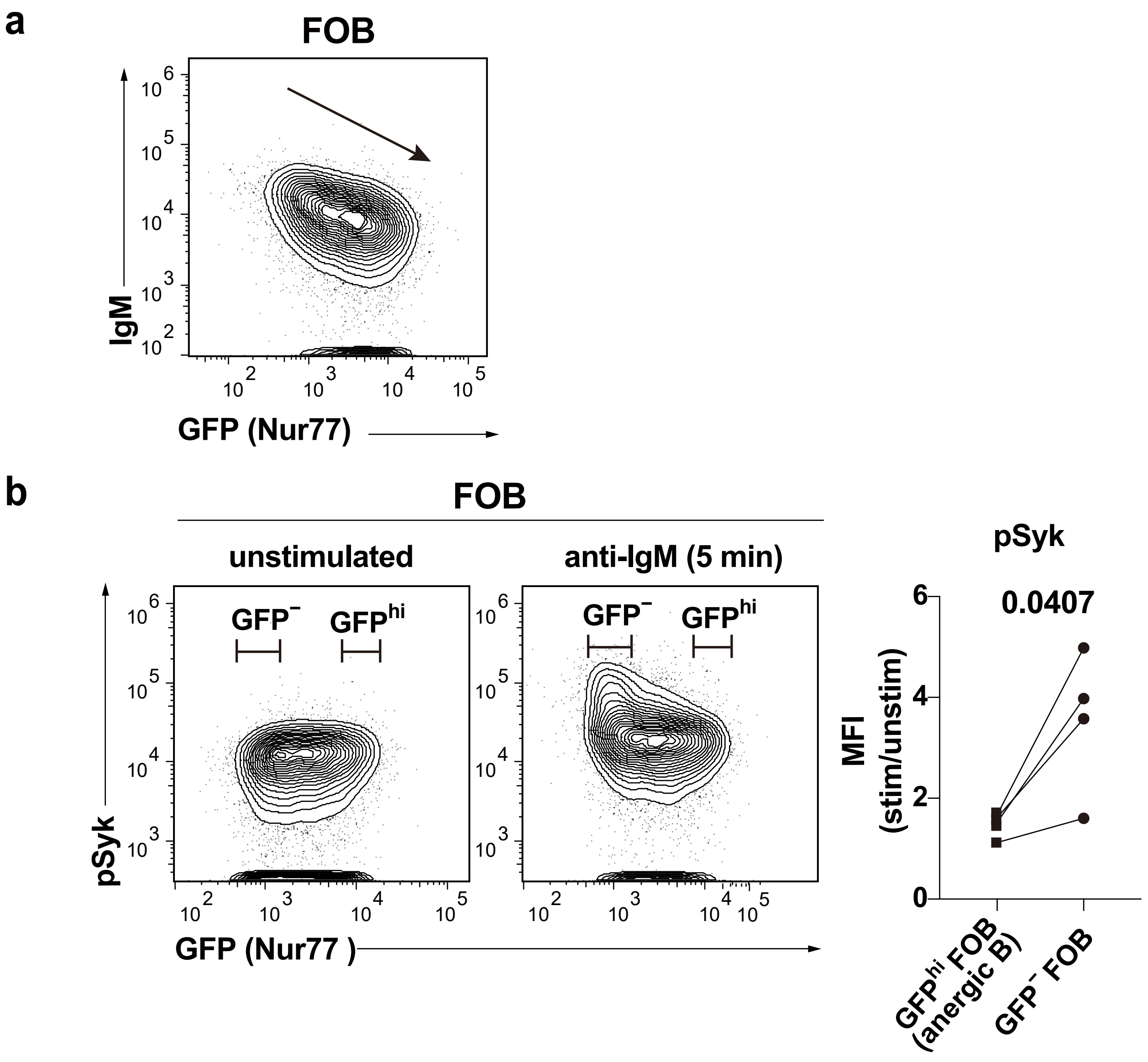


**Extended data Fig. 3 | GFP$^{hi}$ FO B cells derived from Nur77-GFP mice show anergic features. a,** Representative flow cytometry staining of GFP and surface IgM of FO B cells obtained from Nur77 GFP mice (n=5). **b**, Representative flow cytometry staining of GFP and pSyk of splenic FO B cells with or without stimulation with anti-IgM for 5 min. GFP$^{-}$ and GFP$^{hi}$ FO B cells were gated, and the elevated ratio of pSyk (MFI) was compared between GFP$^{-}$ and GFP$^{hi}$ FO B cells. Data are mean ± s.e.m. *P* values are from two-tailed paired Student's *t*-test **(b)**. Data are representative of two **(b)** or three **(a)** independent experiments.

### Extended data Fig. 4

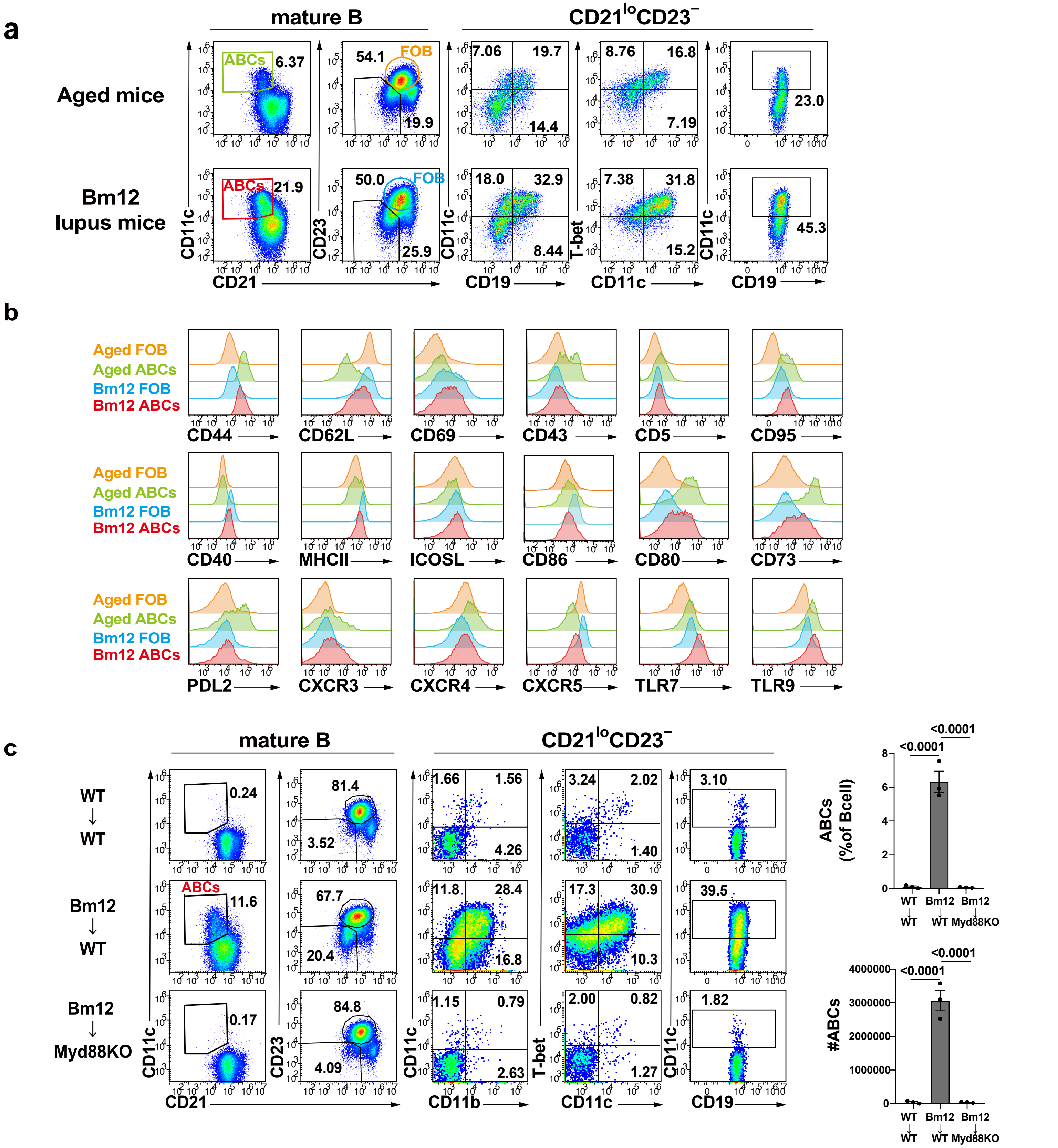


**Extended data Fig. 4 | Characteristics of ABCs in autoimmune mice. a,** Representative flow cytometry plots of splenic FO B cells and ABCs isolated from aged mice (upper, n=3) or Bm12 lupus mice (lower, n=3). **b**, Representative histograms of various markers among splenic FO B cells from aged mice (yellow), ABCs from aged mice (green), FO B cells from Bm12 lupus mice (blue), and ABCs from Bm12 lupus mice (red). **c**, WT mice (n=3) or Myd88KO mice (n=3) were given lymphocytes from Bm12 mice or WT mice and analyzed at day 21. Representative flow cytometry plots of splenic ABCs from WT mice given WT lymphocytes (WT→WT) (n=3), WT mice given Bm12 lymphocytes (Bm12→WT) (n=3), or Myd88KO mice given Bm12 lymphocytes (Bm12→Myd88KO) (n=3) are shown in the left panerel. The percentage and absolute number of splenic ABCs among B cells are shown on the right panel. Data are mean ± s.e.m. *P* values are from one-way ANOVA followed by Tukey's multiple-comparisons test **(c)**: ns, not significant. Data are representative of two independent experiments **(a-c)**.

### Extended data Fig. 5

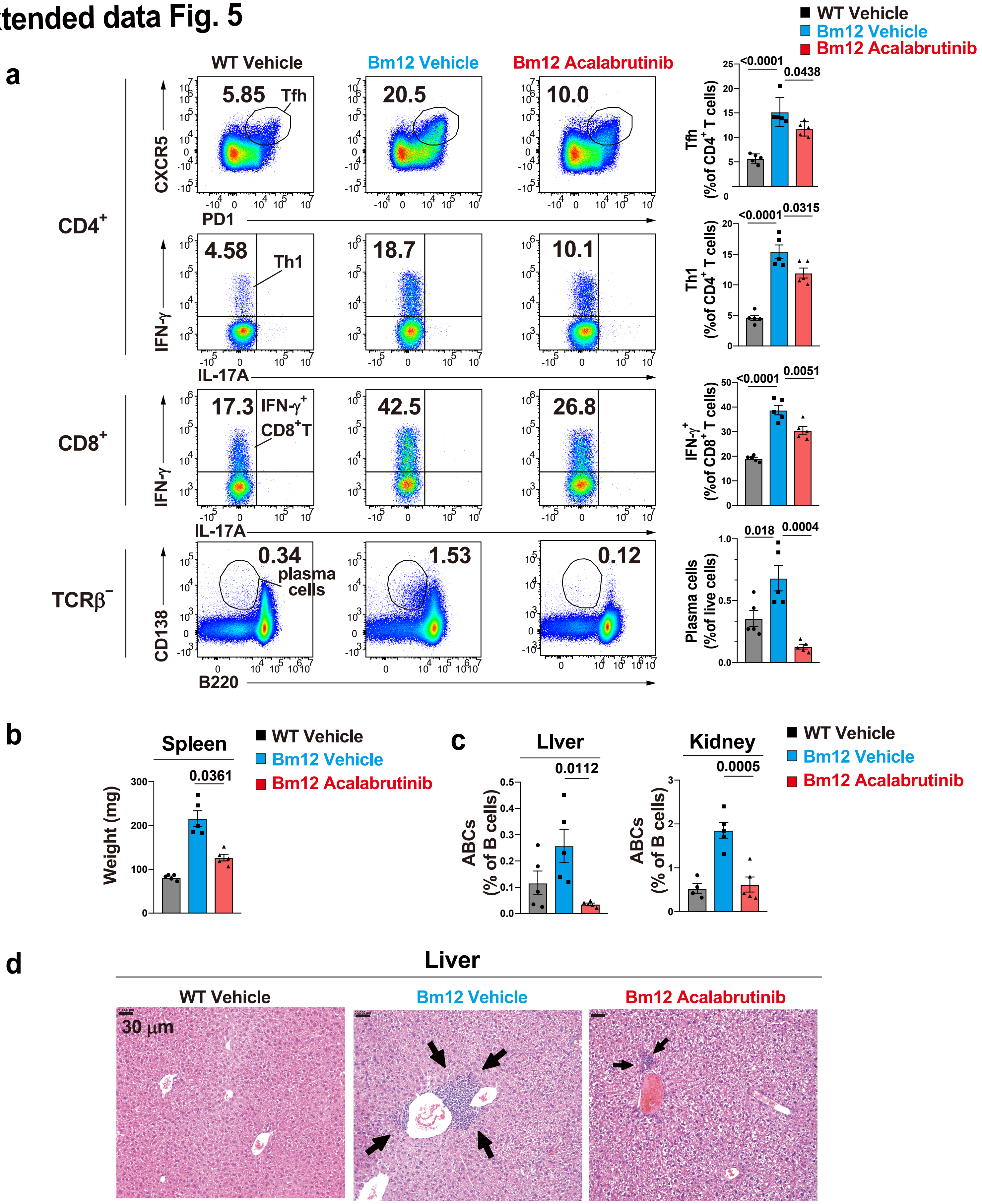


**Extended data Fig. 5 | Btk inhibition ameliorates pathogenesis in lupus-prone mice. a,** Representative flow cytometry plots (left) and the percentage (right) of splenic CD4$^{+}$ Tfh (PD-1$^{hi}$CXCR5$^{hi}$), Th1 (TCRβ$^{+}$CD4$^{+}$IFN-γ$^{+}$), IFN-γ$^{+}$CD8$^{+}$ T cells (TCRβ$^{+}$CD8$^{+}$IFN-γ$^{+}$), and plasma cells (TCRβ$^{-}$B220$^{lo}$CD138$^{+}$) from WT mice given vehicle (n=5, WT Vehicle), Bm12 lupus mice given vehicle (n=5, Bm12 Vehicle), or Bm12 lupus mice given acalabrutinib (n=5, Bm12 Acalabrutinib) in Fig. 3(g). **b**, Spleen weights were compared among these groups. **c**, The percentage of ABCs among B cells collected from liver (left) or kidney (right) were compared among three groups. **d**, Representative H&E-stained histological liver images were compared among three groups. Black arrows show aggregated leukocytes. Scale bars, 30 μm. Data are mean ± s.e.m. *P* values are from two-way ANOVA followed by Tukey's multiple-comparisons test **(a-c)**. Data are representative of two independent experiments **(a-d)**.
